# Developmental dynamic transcriptomics reveals multiple effectors and transcription factors critical for *Ditylenchus destructor* parasitism

**DOI:** 10.1101/2025.06.02.657539

**Authors:** Yangyang Chen, Shurong Zhang, Yayi Zhou, Xueyu Wang, Dexin Bo, Yucheng Liao, Boyan Hu, Yali Zhang, Noha Mohamed Ashry, Donghai Peng, Ming Sun, Dadong Dai

## Abstract

Plant-parasitic nematodes (PPNs) pose a major threat to global agricultural production, yet fundamental research on their biology remains limited. The origin and evolutionary trajectory of PPNs remain elusive, largely due to the scarcity of chromosome-level genomic data. Among them, migratory PPNs are considered a key transitional form between free-living and obligate parasitic lifestyles, as they exhibit both plant parasitism and fungal feeding behaviors. In this study, we assembled a chromosome-level genome of the sweet potato rot nematode *Ditylenchus destructor* and confirmed the presence of four chromosomes through Hi-C scaffolding and karyotype analysis. Comparative genomic analysis with two other migratory PPNs, *Bursaphelenchus xylophilus* and *Aphelenchoides besseyi*, revealed that the Nigon elements in *B. xylophilus* are largely conserved with those of the model organism *Caenorhabditis elegans*, while *D. destructor* and *A. besseyi* exhibit extensive Nigon element rearrangements. These rearrangements were strongly correlated with patterns of protein sequence collinearity. Moreover, transcriptomic profiling across five developmental stages of *D. destructor* identified numerous stage-specific effectors and transcription factors. Functional analysis via RNA interference demonstrated that many of these genes play essential roles in embryogenesis and parasitic activity. Together, our results provide valuable genomic and transcriptomic resources for studying PPNs, uncovering critical insights into their genome evolution and parasitism-related gene functions, and laying a crucial foundation for advancing the understanding of PPN biology and their impact on agricultural systems.

## Introduction

Plant-parasitic nematodes pose a significant threat to global agriculture, causing substantial crop losses and economic burdens[1]. These different types of plant parasitic nematodes can damage the roots, stems, leaves, flowers and fruits of plants, resulting in reduced plant production, a decrease in product quality and yield, as well as co-infection with other pathogens and ultimately plant death[2]. Among them, *Ditylenchus destructor* is a migratory endoparasitic nematode that infects a wide range of host plants, including sweet potatoes and other economically important crops.

Unlike sedentary nematodes, *D. destructor* does not induce and maintain permanent feeding sites and is infectious from larvae to adults[3]. Understanding the genomic and molecular basis of its parasitism is crucial for developing effective nematode management strategies.

The effector of a pathogen is an important weapon for it to infect its host[4]. Previous studies on *D. destructor* have found many effectors that are closely related to parasitism. For example, RNAi of alpha-amylase can reduce the parasitic ability of nematodes by 76.6%[5]; The allergen-like proteins can promote the infection of *D. destructor* nematodes[6]; Dd-cpl-1 RNAi not only leads to a significant decrease in the nematode’s ability to infect, but also causes delayed development of embryos and a decrease in the amount of eggs laid by female[7]. However, no studies have been reported on the function of *D. destructor* transcription factors to date. It is well known that transcription factors regulate the expression of numerous genes and serve as key cis-acting elements in biological processes. For instance, in fungi, the bZIP transcription factor MeaB plays a crucial regulatory role in multiple aspects of Aspergillus fumigatus, including virulence, nitrogen metabolism, biofilm formation, and cell wall integrity[8–10]. Therefore, through high quality genome annotation, exploring the role of these virulence genes and transcription factors in the process of nematode infection of the host will be a meaningful work for analyzing the mechanism of nematode parasitism of the host.

Chromosomal organization and transposable elements (TEs) play pivotal roles in shaping genome architecture, influencing gene expression, and driving genetic diversity[11]. However, the genomic evolution of migratory parasitic nematodes remains largely unexplored, necessitating a detailed investigation into their chromosomal dynamics and TE landscape. Despite the increasing availability of genomic data, the chromosomal evolution of plant-parasitic nematodes remains poorly understood. A key question in nematode genome evolution is how chromosomal rearrangements and transposable elements contribute to adaptation and parasitism. Previous studies have suggested that free-living nematodes, such as *Caenorhabditis elegans*, maintain relatively conserved chromosomal structures, whereas plant-parasitic nematodes exhibit extensive rearrangements[12]. However, the extent of these rearrangements, their impact on gene function, and their evolutionary drivers remain unclear. Furthermore, how these chromosomal changes correlate with parasitic adaptations in different nematode species requires further exploration. Complete genome assemblies are key to advancing studies of chromosome evolution research.

In the early stage of our study, we assembled the genome of *D. destructor* through second-generation sequencing[13], and on this basis, the related functional genes of *D. destructor* were studied[5, 7]. At the same time, we established the transcriptome landscape of 11 time points in *D. destructor* embryonic development[14]. Advancements in sequencing technologies have enabled the construction of high-quality chromosome-level genomes, providing unprecedented insights into nematode evolution and host adaptation. Among this, we utilized third-generation sequencing and chromatin interaction sequencing (Hi-C) to assemble the unzipped genome of the polyploid root-knot nematode, providing novel insights into nematode genomics[12]. In this study, we present a chromosome-level genome assembly of *D. destructor* and perform a comparative genomic analysis with other plant-parasitic nematodes, including *B. xylophilus* and *A. besseyi.* Our long-term goal is to explore fundamental questions regarding genome plasticity and adaptability in plant-parasitic nematodes by investigating chromosome evolution, transposon activity, and the functional roles of secreted proteins and transcription factors in nematode development and parasitism. Our study integrates transcriptomic and RNAi analyses and to provide novel insights into the genetic mechanisms underlying nematode infection, highlighting potential targets for nematode control strategies. This work contributes to a broader understanding of the genomic and evolutionary basis of plant-parasitic nematode adaptations.

## Results

### Chromosome-level genome assembly and annotation of *D. destructor*

The chromosomal evolution of plant-parasitic nematodes remains a relatively understudied field, and the construction of chromosome-level genomes is crucial for advancing research in this area. Firstly, we performed Illumina short-read sequencing on the genomic DNA of *D. destructor* and assessed its ploidy using smudgeplot, which indicated that *D. destructor* is diploid (Fig. S1A) with a haploid genome size of 109Mb (Fig. S1B). To further determine the chromosome number of *D. destructor*, we conducted karyotype analysis, which revealed that *D. destructor* possesses 8 chromosomes (Fig. 1A), i.e., 2n = 8. We utilized 25 GB of PacBio Sequel I data to assemble the genome of *D. destructor*, resulting in a genome size of 105 Mb, an N50 of 899 kb, and a total of 344 contigs. Following polishing with second-generation sequencing data, we generated 40 GB of Hi-C sequencing data. Leveraging the Hi-C data, we successfully anchored 99.92% of the contig sequence length onto four chromosomes (Fig. 1B). Our assembled genome is approximately 30 Mb smaller than the recently published version[15] but has similar BUSCO values (72.2% vs 73.8%).

**Figure 1.**
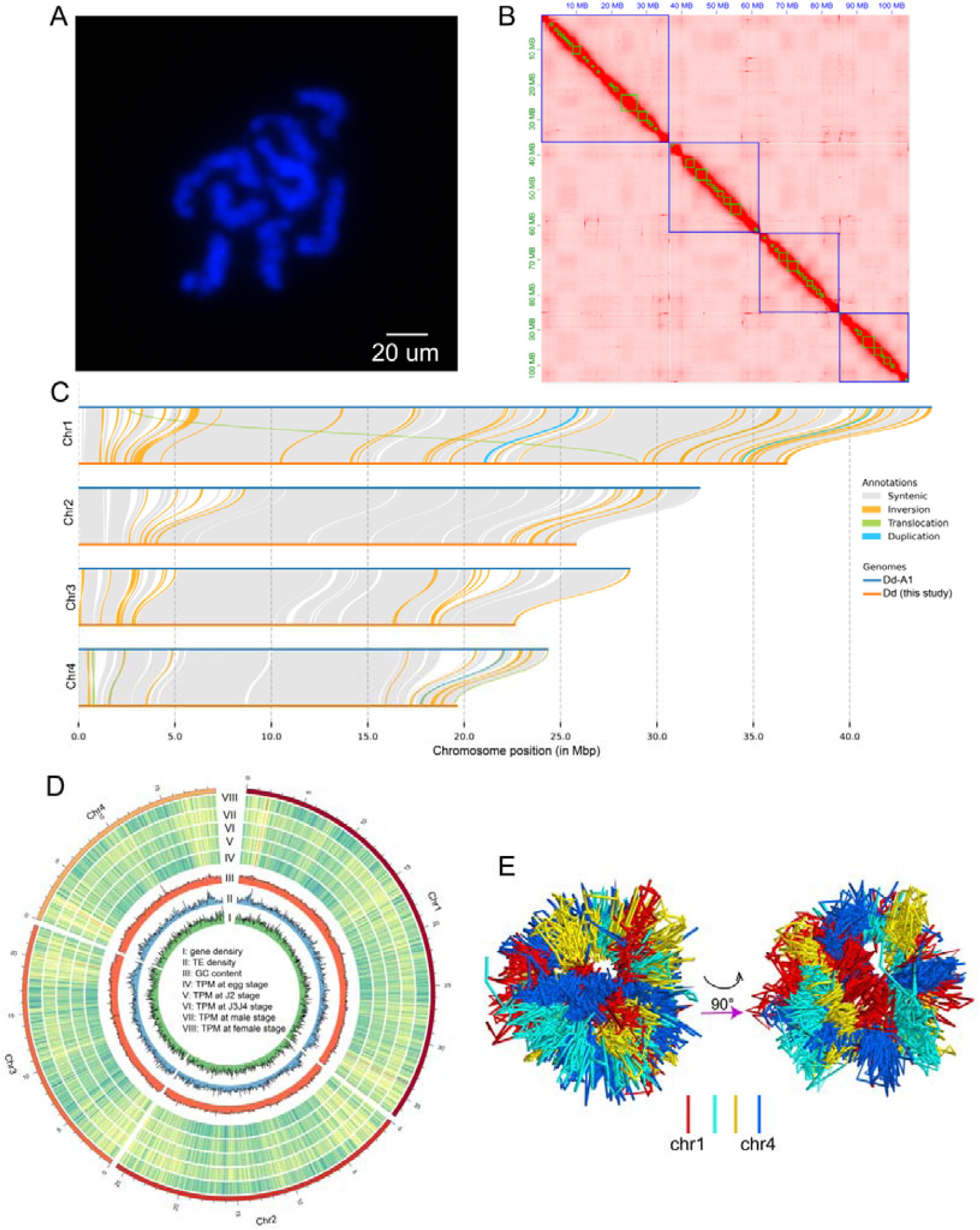
The genome landscape of *D. destructor*. (A) The karyotype photo of *D. destructor* shows that it has 4 pairs of chromosomes. (B) Chromatin interaction signals mount the *D. destructor* genome onto four chromosomes (haploid). (C) Syntenic analysis of *D. destructor* genome between this study and the published Dd-A1 genome (ref14). (D) Circos plot of *D. destructor* haploid genomic features. (E) Reconstructed 3D chromatin structure within the nucleus of *D. destructor* at the mixed developmental stage based on Hi-C data.

Since both the larvae and adults of migratory parasitic nematodes have parasitic abilities, current studies have used transcriptome data from mixed stage to annotate their gene expression. In order to comprehensively study the changes in gene expression of *D. destructor* during development, we used instar synchronization to obtain 18 transcriptome data from six stages: egg, J2, J3J4, male, female, and mixed stage. We identified a total of 20,494 protein coding genes, 563 tRNA, and 48 rRNA in *D. destructor*’s genome. In addition, we annotated 93,855 transposon elements in the genome, with a total length of 23.9 Mb, accounting for 15.9% of the genome size. We then compared our assembled genome with the previously published one[15], highlighting both similarities and differences. Our analysis revealed a high degree of collinearity at the gene level between the two genomes (Fig. S2). Subsequently, we compared the whole genome sequences of the two assemblies and confirmed their high collinearity (Fig. 1C). Additionally, we found that the Dd-A1 genome was larger, with each chromosome containing sequences absent from our assembly (Fig. 1C). In addition, the comparison of the two genomes identified a total of 151,874 SNPs, 26,146 insertions, 23,368 deletions, 93 inversions, 60 translocations, and 42 copygains.

We analyzed various genomic features, including GC content, transposable element (TE) abundance, gene density, and gene expression, and visualized them using circular diagrams (Fig. 1D). To further investigate chromosome organization within the nucleus, we utilized Hi-C data to reconstruct the *D. destructor* chromosome structure and their spatial relationships (Fig. 1E). These findings may offer valuable insights for future studies on gene expression differences and the interplay between age transitions and chromatin dynamics.

### Comparison of genomes among migratory parasitic nematodes

Comparison of chromosomes among plant-parasitic nematodes provides valuable insights into their evolutionary trajectories. We conducted a comparative analysis of chromosome sequences among *D. destructor*, *B. xylophilus* and *A. besseyi*, focusing on their Nigon elements. To facilitate comparison with *Panagrolaimus*, a free-living nematode more closely related to PPNs of the Tylenchomorpha clade[16], we annotated the genome of *Panagrolaimus* sp. ES5. Nigon element analysis revealed that its autosomes are primarily composed of single Nigon elements, whereas the X chromosome represents a fusion of four major elements—NigonA, C, D, and X— suggesting that it originated through fusion events involving two autosomes (Fig. 2A). The Nigon originals of *D. destructor* and *A. besseyi* were fragmented (Fig. 2B, C), suggesting that multiple chromosome divisions and fusions had occurred. To explore the potential association between variations in Nigon elements and chromosomal rearrangements, we performed a synteny analysis of protein sequences across the chromosomes of these four nematode species. Syntenic blocks were identified only when more than ten adjacent protein sequences exhibited similarity across species.

**Figure 2.**
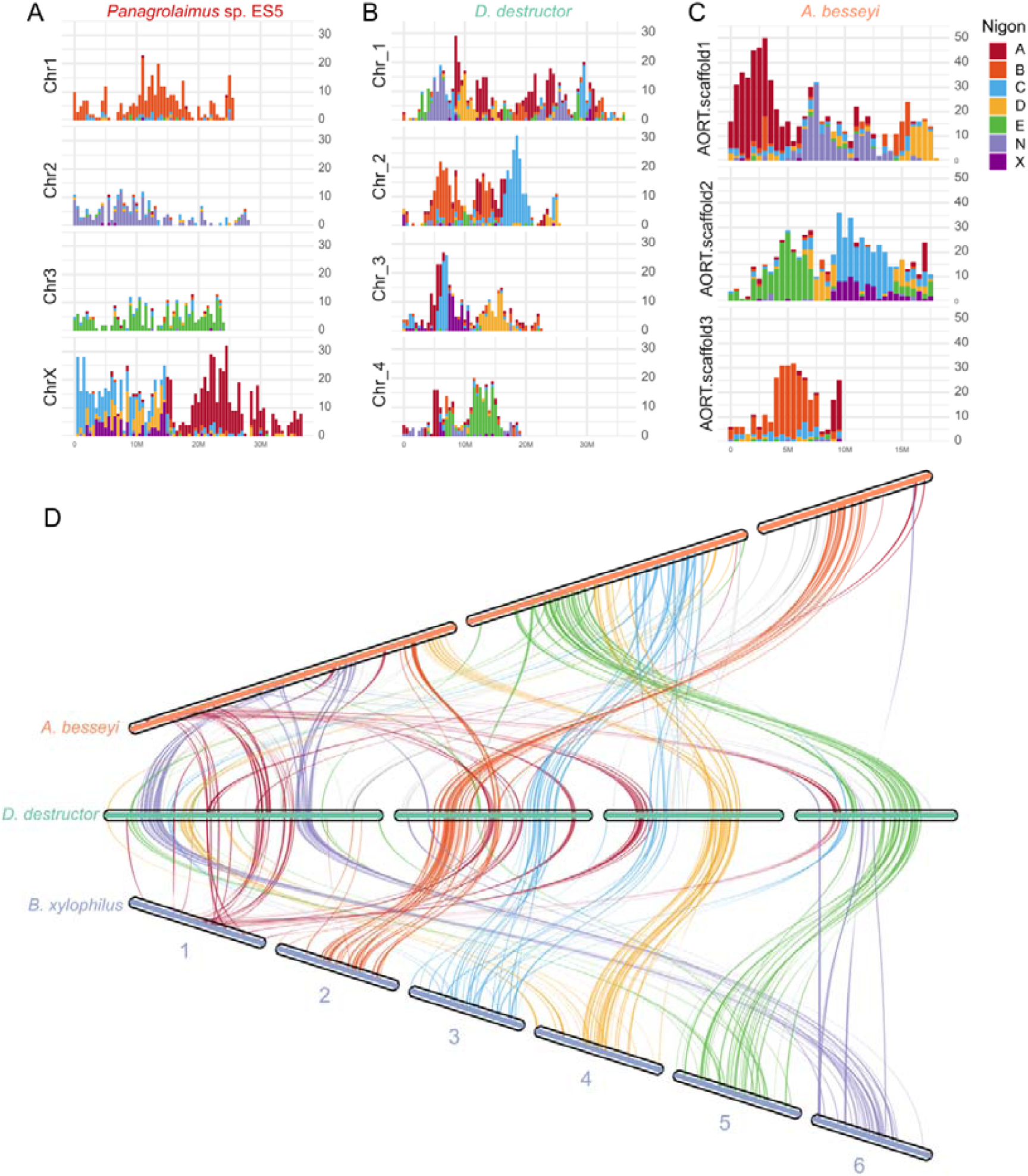
Comparative genomic features of three migratory plant-parasitic nematodes. (A) The distribution characteristics of Nigon elements on the *Panagrolaimus* sp. ES5 genome show that each element is generally concentrated on one autosome. (B) The distribution characteristics of Nigon elements on the *D. destructors* genome. (C) The distribution characteristics of Nigon elements on the *A. besseyi* genome. (D) Synteny of protein sequences among three migratory plant-parasitic nematodes. The results showed that the collinearity of protein sequences was highly consistent with the distribution characteristics of Nigon elements.

Our findings demonstrated a strong concordance between the syntenic blocks of protein sequences and the relative positions of Nigon elements (Fig. 2D). For example, the A Nigon element, which is primarily located on *B. xylophilus* chromosome 1[17], is dispersed across four chromosomes in *D. destructor* but is concentrated on chromosome 1 in *A. besseyi*. This pattern is consistent with the distribution of syntenic blocks and their chromosomal locations. For each of the remaining chromosomes, the positional relationships between Nigon elements and syntenic blocks of protein sequences also exhibit a one-to-one correspondence (Fig. 2). Synteny analysis between *Panagrolaimus* sp. ES5 and *D. destructor* also revealed a high degree of concordance between Nigon element composition and patterns of protein-coding gene collinearity (Fig. S3). Based on this evidence from both Nigon and collinearity, we infer that chromosome 3 of *D. destructor* represents its X chromosome. Our comparative analysis reveals that chromosomal rearrangements, including fusions and fragmentations, have shaped the Nigon structures of PPNs, with *B. xylophilus* retaining a more conserved architecture, while *D. destructor* and *A. besseyi* exhibit extensive chromosomal reorganization.

Transposons elements (TEs) are important drivers of genome dynamics, promoting genome rearrangement, chromosome structural changes, and the evolution of gene regulatory networks[11]. To investigate the role of TEs in migratory plant-parasitic nematodes genome evolution, we conducted a comparative analysis of TEs in these three nematode species. First, we annotated the TE elements in their genomes, revealing that, compared to *B. xylophilus*, *D. destructor* and *A. besseyi* possess a greater diversity of TE families, particularly the LINE transposon family (Fig. S3A, Table S1-3). By analyzing the age distribution of TEs using Kimura substitution rates, we found that the peak age distribution of retrotransposons (LTR/LINE) in *D. destructor* and *A. besseyi* is closer to 0 compared to *B. xylophilus* (Fig. S3B), suggesting that these retrotransposons may have undergone recent active expansion.

### Dynamic landscape of transcriptome at different stages of *D. destructor*

To study the differences in gene expression at different developmental stages of migratory parasitic nematodes, we obtained transcriptome data of *D. destructor* at five stages from egg to adult (Fig. 3A). We first performed differential gene expression analysis between adjacent developmental stages and identified 8,702, 2,881, 4,527, and 4,950 differentially expressed genes (DEGs) in egg_vs_J2, J2_vs_J3J4, J3J4_vs_male, and J3J4_vs_female (Fig. S4), respectively. We conducted a functional analysis of DEGs across developmental stages, providing a valuable resource for identifying key genes involved in plant-parasitic nematode development. The transition from the J3/J4 stage to adult is closely linked to nematode sex determination. Pfam enrichment analysis revealed that genes upregulated in J3/J4 compared to males were primarily associated with protein degradation, particularly ubiquitin ligases such as zf-RING_2, Skp1, Cullin_Nedd8, and CSN8_PSD8_EIF3K (Fig. S5A). Besides, many histone-related genes were enriched in this process, suggesting that sex determination of *D. destructor* may be associated with epigenetics. Moreover, these genes were significantly enriched in essential biological processes, including signal transduction, chromatin regulation, energy metabolism, transcription, DNA replication, and sperm motility (Fig. S5A). Interestingly, the DEGs downregulated in J3/J4 compared to males were primarily enriched in GPCRs, collagen, molting-related genes, and glycosyltransferase families (Fig. S5B). This may suggest that male nematodes enter a relatively stable mature stage, no longer undergoing intense developmental and metabolic remodeling. Instead, they focus more on reproduction while reducing investment in environmental signaling, molting, growth, and stress responses. Surprisingly, the DEGs upregulated or downregulated between J3/J4 and females showed similar functional enrichment patterns to the differential genes between males and females (Fig. S5 C, D). This suggests that J3/J4 larvae share similar genetic regulatory mechanisms during their development into both male and female nematodes. The J3/J4 larval stage appears to be a highly plastic stage, where developmental direction may be controlled by common regulatory networks, with sex determination being an additional fine-tuning of these mechanisms. Furthermore, the differentiation between males and females may rely on fewer key regulatory factors or be influenced by epigenetic regulation, as the DEGs in females did not show enrichment in histone-related genes.

**Figure 3.**
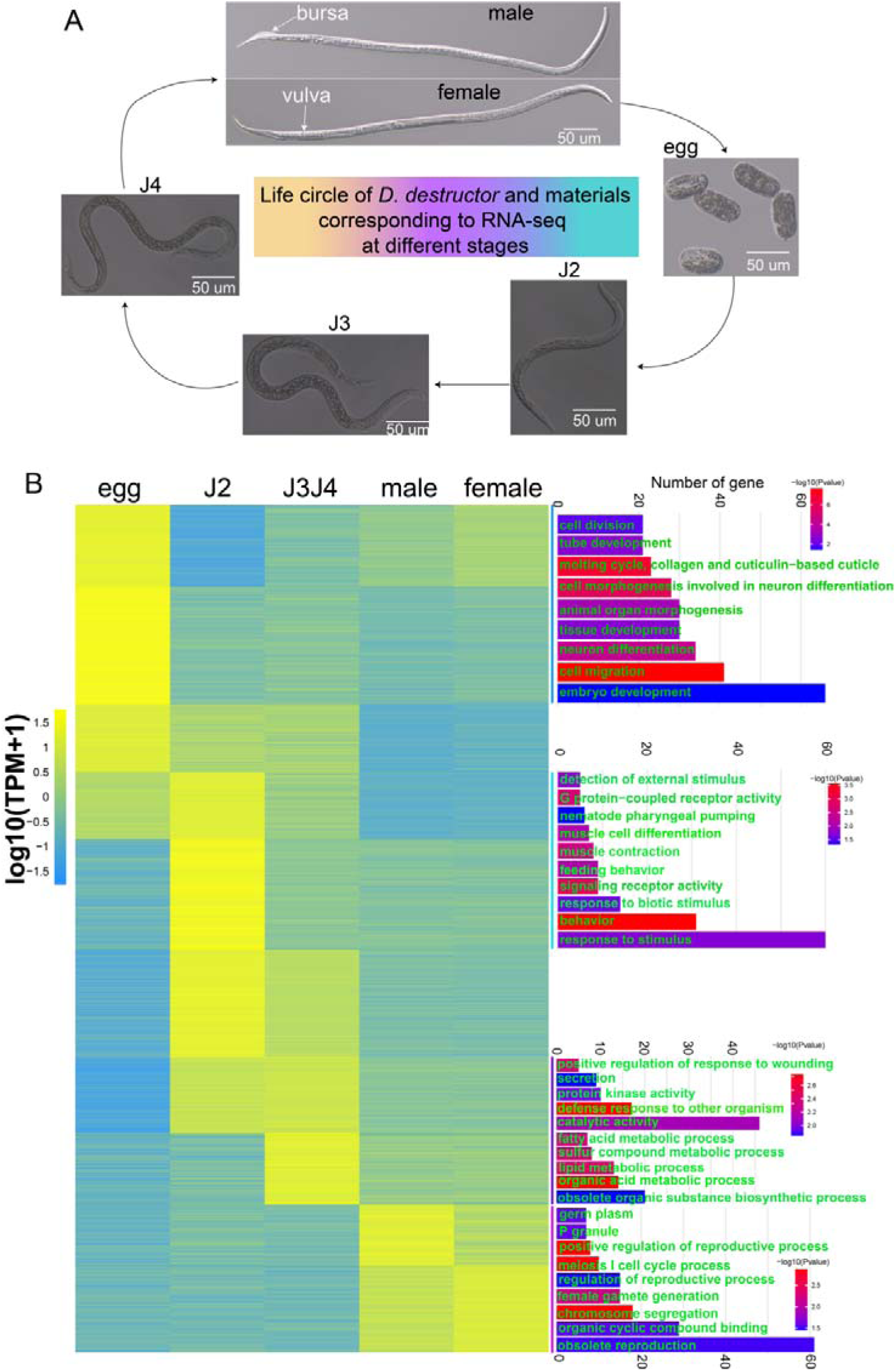
Gene expression profile characteristics and functional analysis of *D. destructor* nematodes at different stages. The heat map on the left shows the clustering of highly expressed genes in each stage after soft clustering of transcriptome data of *D. destructor* at five different developmental stages. The right shows the GO functional enrichment results of genes highly expressed at each developmental stages. The length of the bar plot indicates the number of genes, and the color indicates the p-value of the enrichment degree.

Furthermore, we calculated the Transcripts Per Million (TPM) values of the transcriptomes at each period and performed cluster analysis on the expression levels at different periods. The results indicate that gene expression in *D. destructor* undergoes highly dynamic changes during development (Fig. S6), reflecting finely tuned regulatory mechanisms that adapt to the physiological demands of different developmental stages. To gain deeper insights into gene regulation underlying growth, development, and parasitic strategies, we performed a functional analysis of clustered genes at each developmental stage. The results indicate that the functions of genes enriched at each developmental stage largely correspond to stage-specific characteristics (Fig. 3B). For example, genes highly expressed during the egg stage are primarily enriched in embryonic development, tissue and organ development, and neuronal differentiation. In contrast, genes highly expressed during the J2 to J3/J4 stages are enriched in muscle differentiation, feeding, and responses to environmental stimuli. In the adult stage, enriched functions are related to reproduction, P granules, and gametogenesis (Fig. 3B). These findings suggest that the gene expression patterns of *D. destructor* at different developmental stages are closely linked to their specific physiological functions, reflecting precise gene regulatory mechanisms during development to accommodate the growth demands and survival strategies of each stage.

### Dynamic expression profile of effector, HGT and TF during development

We predicted genes containing secretory signal peptides and no transmembrane domains as secreted proteins, including the nematode pathogenicity factor effector, and annotated 1327 secreted proteins in total. On the other hand, we predicted 124 horizontally transferred genes (HGTs) in the *D. destructor* genome based on heterologous feature analysis. We then analyzed the characteristics of secreted proteins and HGTs in the *D. destructor* genome. In terms of chromosomal distribution, secreted proteins are generally evenly distributed across the chromosomes, with a few regions showing localized clustering (Fig. 4A). In contrast, HGTs exhibit distinct hotspot distributions, particularly concentrated at the chromosomal termini (Fig. 4A). The enrichment of HGTs at chromosome ends may be associated with the dynamic rearrangement, high plasticity, and susceptibility of telomeric regions to foreign gene integration. This distribution pattern may facilitate the retention and adaptive evolution of HGT genes, making them more likely to be preserved by natural selection or further amplified. In addition, we found that 29 of the 124 HGT genes were secreted proteins. For example, all the pectate lyases of *D. destructor* were from HGT, and some of these HGT secreted proteins were concentrated on the chromosomes in hotspot (Fig. 4A).

**Figure 4.**
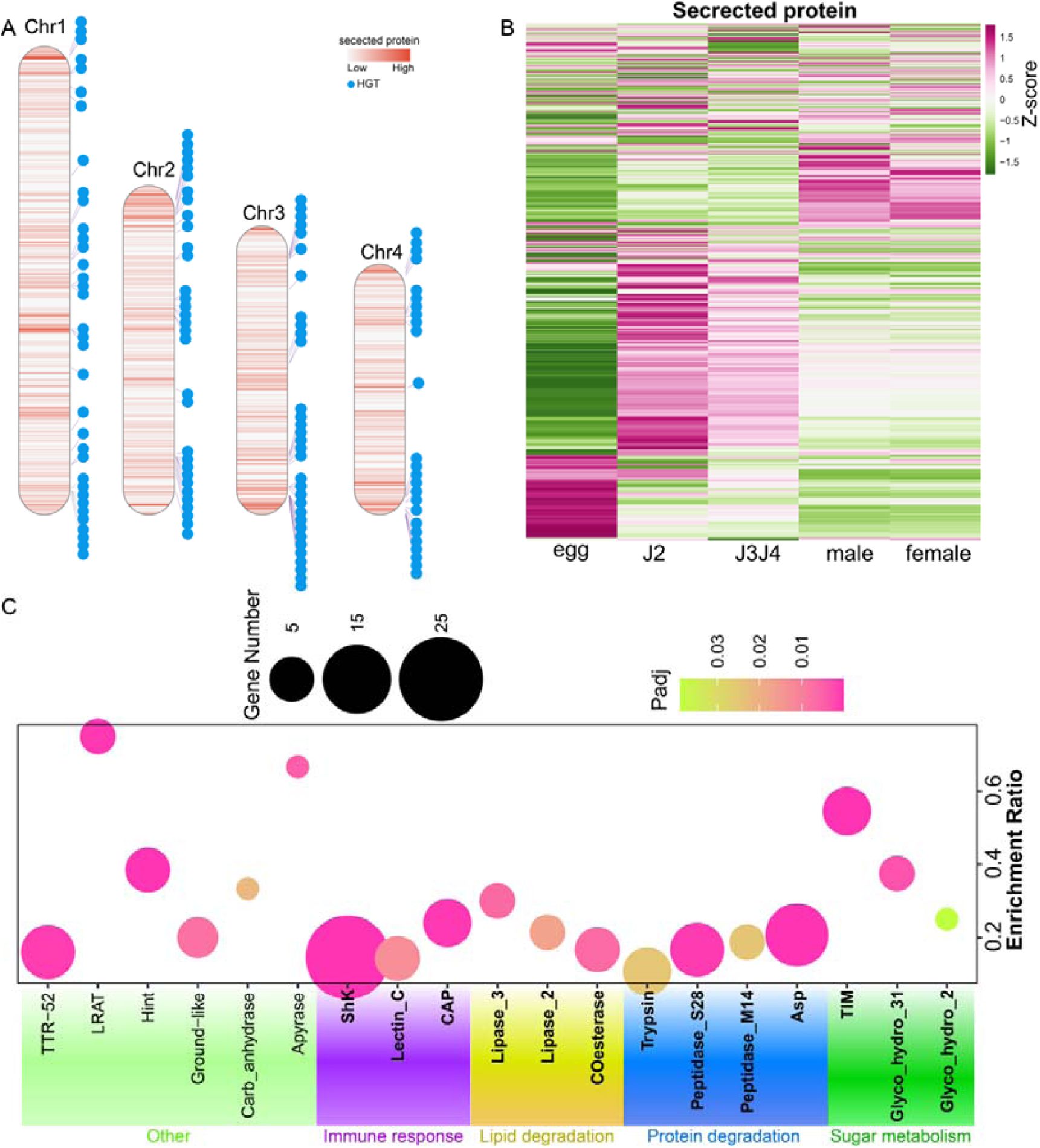
Distribution characteristics of secretory proteins and HGT on the *D. destructor* genome and their expression characteristics among different developmental stages. (A) The distribution characteristics of secretory proteins and HGT on the *D. destructor* genome showed that HGT was distributed in a hotspot pattern, especially concentrated at the chromosome ends. (B) The expression of putative effector (secretory proteins) genes at different developmental ages was clustered, and most of the effectors were highly expressed in the J2 stage. (C) The functional enrichment of putative effectors that were highly expressed in the J2 period was performed through the pfam database. The results showed that a variety of functional genes highly related to nematode feeding, such as those involved in sugar metabolism, protein degradation, and lipid degradation, were enriched.

Although *D. destructor* has the ability to infect at all stages except eggs, we found that most of its secretory proteins have the highest expression levels in the J2 larvae (Fig. 4B), suggesting that the nematode faces many challenges in the initial process of establishing infection. We further conducted functional enrichment analysis on these secretory proteins highly expressed in the J2 period and found that they were mainly enriched in functions such as sugar metabolism, protein degradation, lipid degradation, and immune response (Fig. 4C). These genes play a crucial role in the process of nematodes feeding on host nutrients and resisting immune defense responses, and are the key weapons for nematodes to first infect hosts and establish parasitism. The expression profile of HGT genes during nematode development closely resembles that of secretory proteins, with the majority exhibiting high expression in the J2 larval stage and relatively lower expression in the egg stage (Fig. S7A). In contrast, transcription factors (TFs) display the opposite pattern, with the highest expression observed in the egg stage, followed by the J2 stage (Fig. S7B). This suggests that HGT genes are more closely associated with nematode infection and parasitism, whereas TFs play a crucial regulatory role in embryonic development.

### Secreted proteins and TFs play crucial roles in nematode parasitism

To determine whether these highly expressed secretory proteins and transcription factors (TFs) play a pivotal role in nematode development and host parasitism, we selected four secretory proteins and four TFs that are highly expressed during the J2 stage, along with one TFs that exhibit peak expression during the egg stage (Table S4). RNAi experiments were then conducted to investigate their biological functions (Fig. S8). We first conducted RNAi experiments on the transcription factor gene Dd_05507, which contains a cold shock domain. The results showed that the hatching rate of eggs treated with Dd_05507 dsRNA was significantly lower than that of the control group (Fig. 5A). We then conducted RNAi experiments on secretory protein genes and transcription factors that were highly expressed during the J2 stage. The results revealed that, except for *Dd_09599* and *Dd_03680*, silencing these putative effector genes significantly impaired the nematodes’ ability to infect sweet potatoes (Fig. 5B, C). The RNAi experiments on transcription factors further confirmed that these TFs, belonging to the HMG_box, zf-C4, Homeodomain, and zf-LITAF-like families, are highly expressed during the J2 stage and play a crucial role in facilitating J2 nematodes’ successful colonization of sweet potato and establishment of parasitism. (Fig. 5D, E). These findings highlight the essential role of secretory proteins and transcription factors in nematode development and parasitism. The functional validation of these genes provides key insights into the molecular mechanisms underlying host infection, offering potential targets for disrupting nematode parasitism and developing effective pest control strategies.

**Figure 5.**
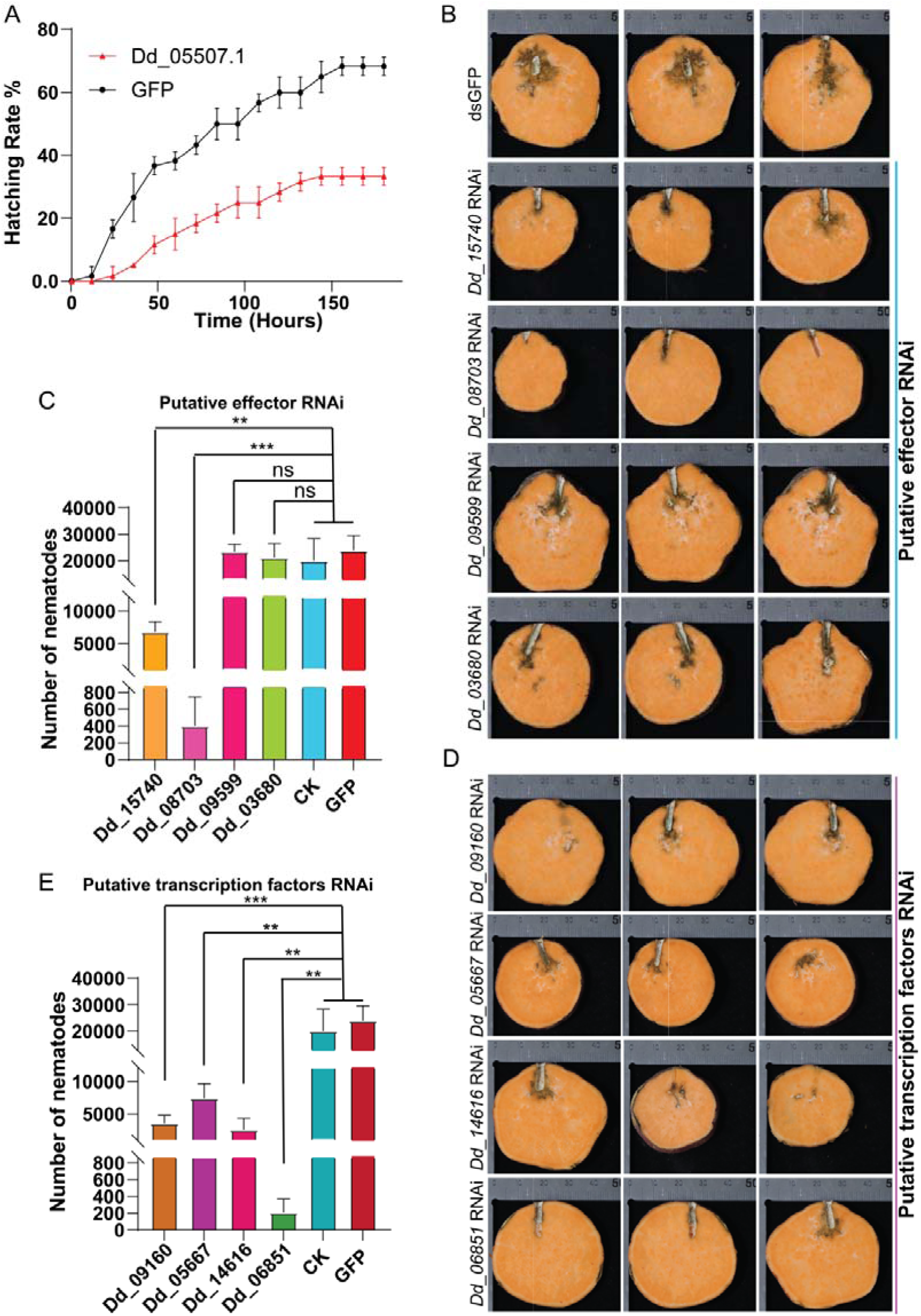
The influence of effector and TF genes in *D. destructor* on its development and infection ability. (A) The hatching rate of eggs was significantly reduced after transcription factor Dd_05507 RNAi. (B) Phenotypes of sweet potato infected with *D. destructor* after the putative effector gene RNAi. (C) Statistics of *D. destructor* reproductive ability after RNAi of putative effector genes. (D) Phenotypes of sweet potato infected with *D. destructor* after the putative TF genes RNAi. (E) Statistics of *D. destructor* reproductive ability after RNAi of putative TF genes. The number of replicates for each treatment in Figures C and E was 4-7, and the significance was analyzed using t-test, ns means no significance, ** means p < 0.01, *** means p < 0.001.

## Disscusion

Recent studies on nematode evolution have shown that *B. xylophilus* is at the root of the phylogenetic tree of plant-parasitic nematodes[16] and has the same chromosome number as *C. elegans*, suggesting that *B. xylophilus* may not have undergone chromosomal recombination, but rather has undergone processes such as inversion, translocation, gene gain and loss, and mutation that enabled it to evolve the ability to parasitize plants. The chromosome-level assembly of *D. destructor* provides a valuable resource for studying the genome evolution of plant-parasitic nematodes.

Our comparative genomic analysis revealed significant chromosomal rearrangements among *D. destructor*, *B. xylophilus*, and *A. besseyi*, with *B. xylophilus* maintaining a more conserved Nigon structure similar to *C. elegans*, whereas *D. destructor* and *A. besseyi* exhibit extensive chromosomal reorganization. These findings suggest that structural variations, including fusions and fragmentations, have played a crucial role in shaping the genome architecture of plant-parasitic nematodes. Transposons elements play an extremely important role in the evolution of many organisms[18–21]. Moreover, TEs can not only regulate genome plasticity, but also act as mediators of genetic variation in plant pathogens, affecting pathogenicity and adaptive evolution[22]. Our transposon analysis indicated that *D. destructor* and *A. besseyi* have undergone recent expansions of retrotransposons, particularly within the LINE family, which may have contributed to genome plasticity and adaptive evolution.

Through years of experience in plant-parasitic nematodes, we have explored a method to synchronize the stages of *D. destructor* and are able to collect nematodes of different stages on a large scale. This work has laid the foundation for detailed studies of the gene function and expression profile of *D. destructor*. The dynamic transcriptomic landscape across developmental stages of *D. destructor* provided insights into the regulatory mechanisms underlying nematode growth and parasitism. Our differential expression analysis revealed stage-specific gene expression patterns, with embryogenesis characterized by genes involved in tissue development and neuronal differentiation, while J2 larvae exhibited upregulation of genes associated with feeding, environmental responses, and host infection. The upregulation of ubiquitin ligases and histone-related genes in J3/J4 suggests potential epigenetic regulation of sex determination, highlighting a complex interplay between chromatin modifications and nematode development.

We identified and characterized secreted proteins and horizontally transferred genes (HGTs) in *D. destructor*, revealing their critical roles in nematode infection and adaptation. Secreted proteins were found to be highly expressed in J2 larvae, underscoring their importance in overcoming host defenses and establishing parasitism. The chromosomal localization of HGTs at telomeric regions suggests their potential involvement in genome plasticity and adaptive evolution[12]. Notably, RNAi-mediated silencing of several highly expressed secreted proteins and significantly impaired nematode infection, confirming their essential roles in parasitism. Additionally, RNAi experiments on transcription factors demonstrated their crucial involvement in J2 colonization and host establishment, providing evidence for the regulatory mechanisms governing nematode development and parasitism. Subsequent work can further explore the mechanism of action between these important effectors and plants and reveal their specific pathogenic processes. At the same time, our results show that TF play a more significant role in nematode colonization. Some transcription factors may regulate the expression of a class of effector proteins or participate in pathways that inhibit plant immunity. Studying the specific regulatory networks of these transcription factors will provide new research ideas for the interaction between PPNs and their hosts.

Our findings provide a comprehensive view of the genetic and regulatory factors that drive nematode parasitism, with implications for developing novel control strategies. The identification of key effectors and transcription factors offers potential targets for disrupting nematode-host interactions, paving the way for more effective pest management approaches.

## Materials and methods

### *D. destructor* nematode culture and collection

Sterilize the surface of sweet potatoes using alcohol and UV in a clean bench for 15 minutes. Punch holes or cut triangular incisions in the sweet potatoes, with an average of two holes per potato, for nematode inoculation. Introduce the counted stem nematodes into the incisions or holes, inoculating 2,000 nematodes per hole. Each sweet potato receives two holes, totaling 4,000 nematodes per potato. Incubate at 25°C with 40%-60% humidity for 30-40 days. Nematodes can then be collected in large quantities using the shallow dish method.

### Instar synchronization of *D. destructor* nematodes

1. **Sample Collection and Processing.** Sweet potatoes inoculated for 30–35 days are collected. The tubers are crushed using a homogenizer at 20% power, and the resulting mixture of nematodes and sweet potato slurry is transferred into a beaker.
2. **Filtration through Sieves.** The mixture is sequentially filtered through a series of standard sieves with mesh sizes of 10, 20, 40, 60, 100, and 200 (determined based on the size of nematodes and eggs), yielding a fine sweet potato particulate fraction mixed with nematodes and eggs.
3. **Density Gradient Centrifugation.** The obtained mixture is aliquoted and subjected to density gradient centrifugation using a 30% sucrose solution to rapidly separate nematodes, eggs, and other impurities. The supernatant, containing nematodes and eggs, is collected and further purified using a 3 µm membrane filter to remove bacteria and debris.
4. **Nematode Lysis and Egg Purification.** The nematode and egg suspension is treated with nematode lysis solution consisting of sodium hypochlorite (50%): sodium hydroxide: water at a ratio of 1:2:7 for 5 minutes, killing the nematodes while preserving viable eggs. Extending the lysis time to 45 minutes ensures complete degradation of nematode bodies, yielding purified, viable eggs.
5. **Egg Hatching and J2 Larval Collection.** The purified eggs are incubated in a 2000-mesh sieve device at 25°C for 72 hours, allowing synchronized hatching of second-stage juveniles (J2).
6. **Isolation of J3 and J4 Juveniles.** The hatched J2 nematodes are inoculated onto Botrytis cinerea plates and incubated at 25°C for 12 days. The nematodes are then collected, and adult nematodes (both male and female) are manually removed. The remaining nematodes consist of third-stage (J3) and fourth-stage (J4) juveniles.
7. **Isolation of male and female.** The female and male insects manually picked in step 6 were saved separately for transcriptome sequencing at the corresponding period.

### DNA, RNA extruction and library construction

DNA samples used for PacBio and Illumina sequencing were obtained from egg stage of *D. destructor*. DNA extraction followed the CTAB method. First, the collected nematodes were centrifuged in a 35% sucrose solution to remove impurities, then suspended in 200 µL of lysis buffer. The nematodes, along with the buffer, were transferred into a mortar containing liquid nitrogen. Once frozen, the nematodes were ground into a fine powder. The ground nematodes were transferred into a 1.5 mL centrifuge tube, followed by the addition of another 300 µL of lysis buffer and 500 µL of preheated (65°C) CTAB. The mixture was thoroughly mixed and incubated at 65°C for 30 minutes, with occasional inversion to ensure proper mixing. Proteins were removed using phenol, RNA was eliminated using RNase A, and DNA was precipitated with isopropanol. DNA intended for PacBio sequencing underwent an additional purification step using magnetic beads to further remove polysaccharide and protein contaminants. For RNA extraction, we first obtained nematodes at different developmental stages using the nematode synchronization method, collecting three biological replicates for each stage. RNA was then extracted using the TransZol Up Plus RNA Kit (TransGen Biotech, Beijinng, China), followed by library construction with the VAHTS Universal V10 RNA-seq Library Prep Kit for Illumina (NR606-01). Finally, RNA sequencing was performed on the Nova platform by NOVOGENE.

### Hi-C library construction

The Hi-C library was constructed using *D. destructor* eggs. The construction method followed the previously established Hi-C library preparation protocol for root-knot nematodes[12]. Briefly, freshly collected eggs were crosslinked with 1.5% formaldehyde for 30 minutes, and the crosslinking was quenched with glycine. The eggs were then ground in a mortar to isolate nuclei. The chromatin was digested with the MboI restriction enzyme, followed by biotin labeling and ligation with T4 DNA ligase. The ligated DNA was then sheared by sonication, and biotin-labeled fragments were enriched using streptavidin magnetic beads. Library preparation was performed using the Vazyme ND606 kit, and the final Hi-C library was sequenced on the Illumina NovaSeq platform with 150 bp paired-end reads.

### Chromosome-level genome assembly

For genome assembly, our workflow was as follows: First, we used Illumina paired-end sequencing data to assess the ploidy of the Dd genome using Smudgeplot[23]. After determining that the genome is diploid, we estimated the genome size using GenomeScope2[23] with a k-mer size of 21, assuming a diploid model. Following genome evaluation, we performed error correction on PacBio Sequel I data using Canu_v1.9_ [24], setting genome coverage at 40×. The corrected 40× data was then assembled using Smartdenovo[25]. The resulting contigs were first polished with gcpp (https://github.com/PacificBiosciences/gcpp) using PacBio data, followed by multiple rounds of polishing with Pilon[26] using Illumina data. Next, we used Juicer (v1.5.7) pipline[27] and 3D-DNA (v180419) pipline[28] to anchor the polished genome to chromosomes based on Hi-C signals. Finally, the anchored genome was manually refined in Juicebox (v1.13.01) [29], yielding the final chromosome-level genome assembly. The three-dimensional structure of the genome was reconstructed using miniMDS[30] software.

### Genome annotation

We used transcriptome data from all developmental stages to assist in annotating the genome of *Ditylenchus destructor*. First, we employed Trinity (v2.15.2)[31] to perform de novo assembly of the transcriptome data, resulting in the “Trinity.fasta” file. The assembled sequences were then used in PASA[32] for ORF prediction, generating the “pasa_assemblies.gff3” file. Next, we masked the Dd genome using RepeatMasker[33], followed by alignment of the transcriptome data to the masked genome with HISAT2[34]. The resulting BAM files were used as input for BRAKER3[35] to predict genes, generating the “braker.gff3” file. Additionally, the BAM files from HISAT2 alignment were used for gene prediction with TransDecoder and StringTie (v2.1.1)[36], yielding the “stringtie.gff3” file. Finally, we merged the gene predictions from the three methods using EVidenceModeler, incorporating weight-based merging to produce the final annotation file, “annotation.gff3”. CDS and protein sequences were extracted from this GFF3 file using GFFRead. The genome annotation of *Panagrolaimus* sp. ES5 was performed by using the same strategy, the genome and RNA-seq data were download from NCBI with accession number GCA_964243005.1 and SRR5253562.

### TE, HGT, TF, and secretory protein prediction

For TE prediction, we first used EDTA (v2.2.0) pipline[37] to predict the genome with the parameters “--overwrite 1 --force 1 --sensitive 1 --anno 1 --evaluate 1” and obtained “TEanno.gff3” and “TElib.fa”. Then, using “TElib.fa” as a library file, RepeatMasker[33] was used to re-predict TE in the genome, and calcDivergenceFromAlign.pl was used to calculate Kimura substitution on the result file "*.align". For HGT prediction, we use AvP pipline[38] with uniref90 database, the candidate gene with HGT or HGT-NT lable was considered as HGT genes. For transcription factors, we divided all protein sequences of *D. destructor* into two groups of no more than 9000 genes and predict the TF by using AnimalTFDB v4.0 (https://guolab.wchscu.cn/AnimalTFDB4/#/TF_Predict). For secretory protein prediction, we first used singalP(v5.0)[39] to predict genes with secretory signal peptides, and then used tmhmm[40] to predict whether these genes had transmembrane domains. After eliminating genes containing transmembrane domains, they were potential secretory proteins.

### Collinearity and genomic variation analysis

We used minimap2 with the parameters “-ax asm5 --eqx” to align the entire genome between the *D. destructor* genome of this study and Dd-A1[15], and then used Syri[41] to identify collinear regions, structural variations, insertion and deletion mutations, and SNP information. The syntenic protein pairs between the *D. destructor* genome of this study and Dd-A1 were identified by using JCVI pipeline[42].

### RNA-seq analysis and expression profile determination

For the analysis of differentially expressed genes, we first aligned the transcriptome data to the genome using HISAT2, followed by sorting with SAMtools[43] and quantifying read counts with HTSeq[44]. Differential gene expression analysis was then performed using DESeq2[45]. For TPM calculation, we first aligned the transcriptome data to the genome using STAR[46] and then used RSEM[47] to calculate TPM values for each gene. Gene expression profile clustering across developmental stages was conducted using Mfuzz[48], while the expression pattern clustering of transcription factors, effectors, and other key genes at different stages was performed using pheatmap.

### In vitro RNAi

Primers were designed for target genes, each incorporating a T7 promoter sequence. Then, the target genes were amplified from *D. destructor* cDNA. The PCR products were purified, ensuring DNA concentrations exceeded 100 ng/µl. Vazyme T7 RNAi Transcription Kit (TR102) was used for dsRNA synthesis. The final dsRNA concentration was adjusted to 1000 ng/µl. For the RNAi interference system preparation, the total reaction volume was 300 µl, supplemented with 15 µl of 1 mol/L octopamine and 0.5 µL of 1 mol/L spermidine. For the nematode and egg RNAi treatment, we use 6000 nematodes or 5000 eggs were for each gene-specific RNAi system. The interference reaction was conducted at 37°C for 48 hours. After RNAi treatment, 2000 nematodes were collected for RNA extraction and then used for qRT-PCR, the remaining nematodes were used for phenotypic analysis.

### Assessment of nematode infectivity

Infection assay: Sweet potatoes of uniform diameter were punctured to a standardized depth (about 1cm). Each hole was inoculated with 500 nematodes, with 8 replicates per gene. After 30 days infection at 25°C of dark incubator, nematodes were collected and counted. The extent of infection was examined, and photographs were taken for ocumentation.

Egg hatching rate assay: 30–40 early-stage embryos were selected per treatment, with 5 replicates per gene. The eggs were incubated at 25°C for 3–7 days, hatching rates were recorded every 24 hours.

Nematode quantification: Harvesting nematodes from infected sweet potatoes, a 2 cm region surrounding the infection site was excised. The infected tissue was cut into 0.5 cm × 0.5 cm pieces and wrapped in a single layer of paper towel. The samples were submerged in water for 4 hours to allow nematodes to migrate into the liquid. Nematode suspensions were collected and counted at least three times to estimate the total population.

Negative Control Group: dsGFP synthesized using GFP gene as a template was used as a negative control for RNAi. The same infection ability and egg hatching rate assays were performed for comparison.

## Data availability

All sequencing data have been deposited into the National Center for Biotechnology Information (NCBI) Sequence Read Archive. The different stages of RNA sequencing data of *D. destructor* are deposited in BioProject PRJNA1164248 (including 15 SRA); The PacBio Sequel I data, Hi-C data, and Illumina data of *D. destructor* are deposited in BioProject PRJNA1236705 (including 3 SRA). The genome of *B. xylophilus* and *A. besseyi* were download form wormbase parasite with accession number PRJEB40022 and PRJNA834627. The genome of *Panagrolaimus* sp. ES5 was download from GenBank with accession number GCA_964243005.1, and the RNA-seq data was download from SRA database with accession number SRR5253562. The assembled *D. destructor* genome and annotation files were uploaded to https://bmb.hzau.edu.cn/sjxz.htm.

## Acknowledgements

We thank Romnick Latina of University of California Davis for advice on nematodes evolution and the UC Davis farm cluster server for help. This research was supported by the National Natural Science Foundation of China (32271546, U20A2040 and 31970076) and Hubei Hongshan Laboratory (2022hszd012).

## Author Contributions

D.D. and M.S. conceived and administration the project. M.S., acquisition the funding. M.S. and D.P. manage this project. D.D. designed the experiments. Y.C. collected the data and performed the RNAi experiment. S. Z. clone the gene. S. Z. and X.W. constructed the RNA-seq libraries. D.D. and Y.Z. performed the data analysis. B.H., D.B., Y.L., N.A., and Y.L.Z. raised nematodes. D.D. wrote the manuscript. All authors reviewed the manuscript.

## References

1. Jones JT, Haegeman A, Danchin EG, Gaur HS, Helder J, Jones MG, Kikuchi T, Manzanilla-Lopez R, Palomares-Rius JE, Wesemael WM et al: Top 10 plant-parasitic nematodes in molecular plant pathology. Mol Plant Pathol 2013, 14(9):946–961.

2. Parrado LM, Quintanilla M: Plant-parasitic nematode disease complexes as overlooked challenges to crop production. Front Plant Sci 2024, 15:1439951.

3. Lu Y, Yang S, Chen W, Xie H, Xu C: Advances in Migratory Plant Endoparasitic Nematode Effectors. Int J Mol Sci 2024, 25(12).

4. Lanver D, Tollot M, Schweizer G, Lo Presti L, Reissmann S, Ma LS, Schuster M, Tanaka S, Liang L, Ludwig N et al: Ustilago maydis effectors and their impact on virulence. Nat Rev Microbiol 2017, 15(7):409–421.

5. Chen L, Xu M, Wang C, Zheng J, Huang G, Chen F, Peng D, Sun M: Multi-copy alpha-amylase genes are crucial for Ditylenchus destructor to parasitize the plant host. PLoS One 2020, 15(10):e0240805.

6. Chang Q, Yang Y, Hong B, Zhao Y, Zhao M, Han S, Zhang F, Peng H, Peng D, Li Y: A variant of the venom allergen-like protein, Dd VAP2, is required for the migratory endoparasitic plant nematode Ditylenchus destructor parasitism of plants. Frontiers in Plant Science 2023, 14:1322902.

7. Huang G, Cong Z, Liu Z, Chen F, Bravo A, Soberon M, Zheng J, Peng D, Sun M: Silencing Ditylenchus destructor cathepsin L-like cysteine protease has negative pleiotropic effect on nematode ontogenesis. Sci Rep 2024, 14(1):10030.

8. Wagner D, Schmeinck A, Mos M, Morozov IY, Caddick MX, Tudzynski B: The bZIP transcription factor MeaB mediates nitrogen metabolite repression at specific loci. Eukaryotic Cell 2010, 9(10):1588–1601.

9. Perez-Cuesta U, Guruceaga X, Cendon-Sanchez S, Pelegri-Martinez E, Hernando FL, Ramirez-Garcia A, Abad-Diaz-de-Cerio A, Rementeria A: Nitrogen, iron, and zinc acquisition: key nutrients to Aspergillus fumigatus virulence. Journal of Fungi 2021, 7(7):518.

10. Chen Y, Gao F, Chen X, Tao S, Chen P, Lin W: The basic leucine zipper transcription factor MeaB is critical for biofilm formation, cell wall integrity, and virulence in Aspergillus fumigatus. Msphere 2024, 9(2):e00619–00623.

11. Slotkin RK, Martienssen R: Transposable elements and the epigenetic regulation of the genome. Nat Rev Genet 2007, 8(4):272–285.

12. Dai D, Xie C, Zhou Y, Bo D, Zhang S, Mao S, Liao Y, Cui S, Zhu Z, Wang X et al: Unzipped chromosome-level genomes reveal allopolyploid nematode origin pattern as unreduced gamete hybridization. Nat Commun 2023, 14(1):7156.

13. Zheng J, Peng D, Chen L, Liu H, Chen F, Xu M, Ju S, Ruan L, Sun M: The Ditylenchus destructor genome provides new insights into the evolution of plant parasitic nematodes. Proc Biol Sci 2016, 283(1835).

14. Wang X, Guo Z, Dai D, Xie C, Zhao Z, Zheng J, Sun M, Peng D: High-resolution transcriptome datasets during embryogenesis of plant-parasitic nematodes. Sci Data 2024, 11(1):690.

15. Yang Y, Feng R, Hong B, Fang Y, Liu C, Wang K, Peng D, Li Y, Peng H, Chang Q: Chromosome-level genome assembly of the sweet potato rot nematode Ditylenchus destructor. Sci Data 2025, 12(1):174.

16. Qing X, Zhang YM, Sun S, Ahmed M, Lo WS, Bert W, Holovachov O, Li H: Phylogenomic Insights into the Evolution and Origin of Nematoda. Syst Biol 2024.

17. Gonzalez de la Rosa PM, Thomson M, Trivedi U, Tracey A, Tandonnet S, Blaxter M: A telomere-to-telomere assembly of Oscheius tipulae and the evolution of rhabditid nematode chromosomes. *G3* (*Bethesda*) 2021, 11(1).

18. Wicker T, Gundlach H, Spannagl M, Uauy C, Borrill P, Ramirez-Gonzalez RH, De Oliveira R, International Wheat Genome Sequencing C, Mayer KFX, Paux E et al: Impact of transposable elements on genome structure and evolution in bread wheat. Genome Biol 2018, 19(1):103.

19. Kumar CS, Qureshi SF, Ali A, Satyanarayana ML, Rangaraju A, Venkateshwari A, Nallari P: Hidden magicians of genome evolution. Indian J Med Res 2013, 137(6):1052–1060.

20. Yu Z, Li J, Wang H, Ping B, Li X, Liu Z, Guo B, Yu Q, Zou Y, Sun Y et al: Transposable elements in Rosaceae: insights into genome evolution, expression dynamics, and syntenic gene regulation. Hortic Res 2024, 11(6):uhae118.

21. Shao F, Han M, Peng Z: Evolution and diversity of transposable elements in fish genomes. Sci Rep 2019, 9(1):15399.

22. Mat Razali N, Cheah BH, Nadarajah K: Transposable Elements Adaptive Role in Genome Plasticity, Pathogenicity and Evolution in Fungal Phytopathogens. Int J Mol Sci 2019, 20(14).

23. Ranallo-Benavidez TR, Jaron KS, Schatz MC: GenomeScope 2.0 and Smudgeplot for reference-free profiling of polyploid genomes. Nat Commun 2020, 11(1):1432.

24. Koren S, Walenz BP, Berlin K, Miller JR, Bergman NH, Phillippy AM: Canu: scalable and accurate long-read assembly via adaptive k-mer weighting and repeat separation. Genome Res 2017, 27(5):722–736.

25. Liu H, Wu S, Li A, Ruan J: SMARTdenovo: a de novo assembler using long noisy reads. GigaByte 2021, 2021:gigabyte15.

26. Walker BJ, Abeel T, Shea T, Priest M, Abouelliel A, Sakthikumar S, Cuomo CA, Zeng Q, Wortman J, Young SK et al: Pilon: an integrated tool for comprehensive microbial variant detection and genome assembly improvement. PLoS One 2014, 9(11):e112963.

27. Durand NC, Shamim MS, Machol I, Rao SS, Huntley MH, Lander ES, Aiden EL: Juicer Provides a One-Click System for Analyzing Loop-Resolution Hi-C Experiments. Cell Syst 2016, 3(1):95–98.

28. Dudchenko O, Batra SS, Omer AD, Nyquist SK, Hoeger M, Durand NC, Shamim MS, Machol I, Lander ES, Aiden AP et al: De novo assembly of the Aedes aegypti genome using Hi-C yields chromosome-length scaffolds. Science 2017, 356(6333):92-95.

29. Durand NC, Robinson JT, Shamim MS, Machol I, Mesirov JP, Lander ES, Aiden EL: Juicebox Provides a Visualization System for Hi-C Contact Maps with Unlimited Zoom. Cell Syst 2016, 3(1):99–101.

30. Rieber L, Mahony S: miniMDS: 3D structural inference from high-resolution Hi-C data. Bioinformatics 2017, 33(14):i261–i266.

31. Grabherr MG, Haas BJ, Yassour M, Levin JZ, Thompson DA, Amit I, Adiconis X, Fan L, Raychowdhury R, Zeng Q: Trinity: reconstructing a full-length transcriptome without a genome from RNA-Seq data. Nature biotechnology 2011, 29(7):644.

32. Haas BJ, Salzberg SL, Zhu W, Pertea M, Allen JE, Orvis J, White O, Buell CR, Wortman JR: Automated eukaryotic gene structure annotation using EVidenceModeler and the Program to Assemble Spliced Alignments. Genome Biol 2008, 9(1):R7.

33. Tarailo-Graovac M, Chen N: Using RepeatMasker to identify repetitive elements in genomic sequences. Curr Protoc Bioinformatics 2009, Chapter 4:4 10 11-14 10 14.

34. Kim D, Paggi JM, Park C, Bennett C, Salzberg SL: Graph-based genome alignment and genotyping with HISAT2 and HISAT-genotype. Nat Biotechnol 2019, 37(8):907–915.

35. Gabriel L, Bruna T, Hoff KJ, Ebel M, Lomsadze A, Borodovsky M, Stanke M: BRAKER3: Fully automated genome annotation using RNA-seq and protein evidence with GeneMark-ETP, AUGUSTUS, and TSEBRA. Genome Res 2024, 34(5):769–777.

36. Pertea M, Pertea GM, Antonescu CM, Chang TC, Mendell JT, Salzberg SL: StringTie enables improved reconstruction of a transcriptome from RNA-seq reads. Nat Biotechnol 2015, 33(3):290–295.

37. Ou S, Su W, Liao Y, Chougule K, Agda JRA, Hellinga AJ, Lugo CSB, Elliott TA, Ware D, Peterson T et al: Benchmarking transposable element annotation methods for creation of a streamlined, comprehensive pipeline. Genome Biol 2019, 20(1):275.

38. Koutsovoulos GD, Granjeon Noriot S, Bailly-Bechet M, Danchin EGJ, Rancurel C: AvP: A software package for automatic phylogenetic detection of candidate horizontal gene transfers. PLoS Comput Biol 2022, 18(11):e1010686.

39. Almagro Armenteros JJ, Tsirigos KD, Sonderby CK, Petersen TN, Winther O, Brunak S, von Heijne G, Nielsen H: SignalP 5.0 improves signal peptide predictions using deep neural networks. Nat Biotechnol 2019, 37(4):420–423.

40. Krogh A, Larsson B, von Heijne G, Sonnhammer EL: Predicting transmembrane protein topology with a hidden Markov model: application to complete genomes. J Mol Biol 2001, 305(3):567–580.

41. Goel M, Sun H, Jiao WB, Schneeberger K: SyRI: finding genomic rearrangements and local sequence differences from whole-genome assemblies. Genome Biol 2019, 20(1):277.

42. Wang Y, Tang H, Debarry JD, Tan X, Li J, Wang X, Lee TH, Jin H, Marler B, Guo H et al: MCScanX: a toolkit for detection and evolutionary analysis of gene synteny and collinearity. Nucleic Acids Res 2012, 40(7):e49.

43. Li H, Handsaker B, Wysoker A, Fennell T, Ruan J, Homer N, Marth G, Abecasis G, Durbin R, Genome Project Data Processing S: The Sequence Alignment/Map format and SAMtools. Bioinformatics 2009, 25(16):2078–2079.

44. Anders S, Pyl PT, Huber W: HTSeq--a Python framework to work with high-throughput sequencing data. Bioinformatics 2015, 31(2):166–169.

45. Love MI, Huber W, Anders S: Moderated estimation of fold change and dispersion for RNA-seq data with DESeq2. Genome Biol 2014, 15(12):550.

46. Dobin A, Davis CA, Schlesinger F, Drenkow J, Zaleski C, Jha S, Batut P, Chaisson M, Gingeras TR: STAR: ultrafast universal RNA-seq aligner. Bioinformatics 2013, 29(1):15–21.

47. Li B, Dewey CN: RSEM: accurate transcript quantification from RNA-Seq data with or without a reference genome. BMC Bioinformatics 2011, 12:323.

48. Kumar L, M EF: Mfuzz: a software package for soft clustering of microarray data. Bioinformation 2007, 2(1):5–7.

