## Supplemental Figures for "Developmental dynamic transcriptomics reveals multiple effectors and transcription factors critical for *Ditylenchus destructor* parasitism"

### Supplementary Figures

**Fig. S1.** Ploidy and genome size estimation of D. destructor.
**Fig. S2.** Colinearity analysis of gene sequences between the *D. destructor* genome in this study and the published Dd-A1 genome.
**Fig. S3.** The TE landscape of three migratory plant-parasitic nematodes.
**Fig. S4.** Volcano plot showing differentially expressed genes between different developmental stages of *D. destructor*.
**Fig. S5.** Functional enrichment analysis of differentially expressed genes between J3J4 larvae and adults.
**Fig. S6.** Clustering of TPM values of gene expression between different stages of *D. destructor* shows age-specific expression profiles.
**Fig. S7.** The expression pheatmap of HGT and TF genes during the development stage.
**Fig. S8.** qRT-PCR analysis of the relative expression levels of the candidate effector gene and transcription factor target genes following RNAi treatment.

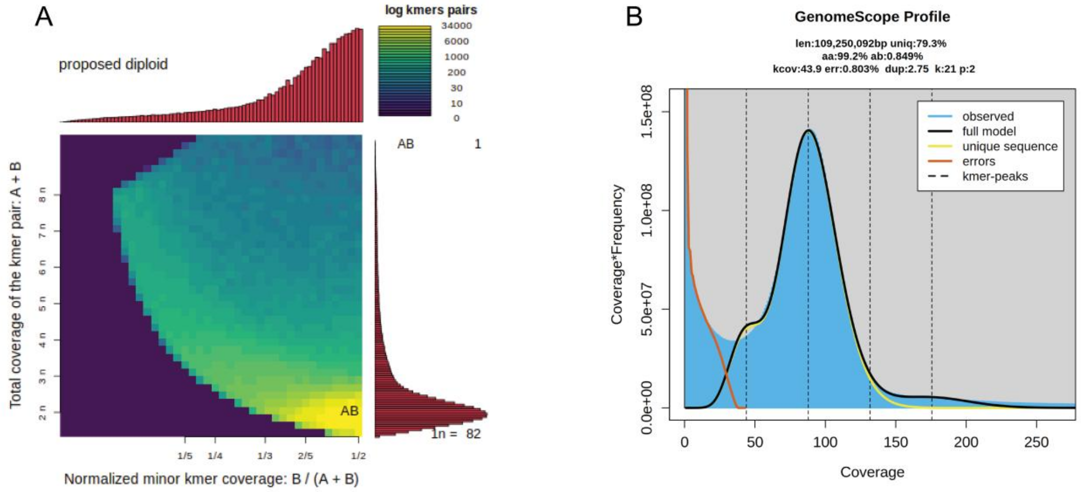


**Fig. S1.** Ploidy and genome size estimation of D. destructor. **A:** Smudgeplot evaluates that the genome of Dd is diploid. **B:** GenomeScope2 estimated the genome size of Dd to be 109 Mb.

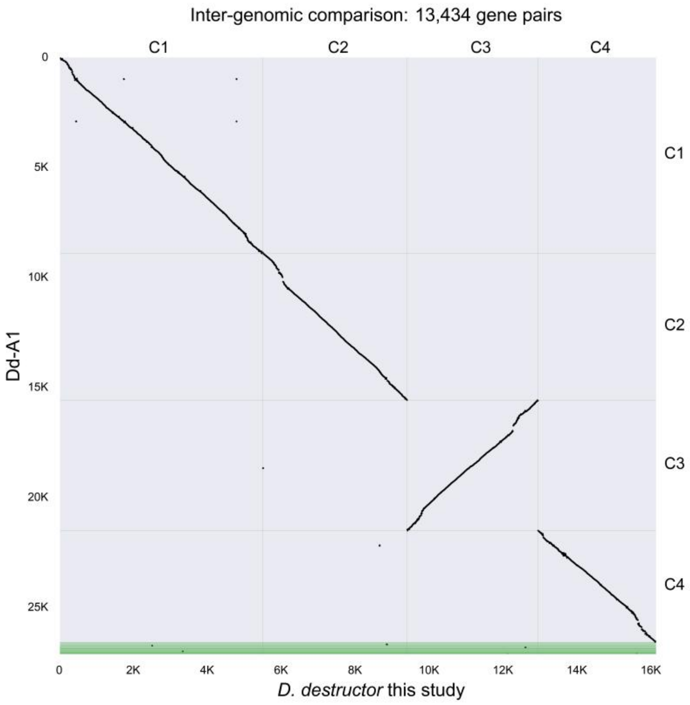


**Fig. S2.** Colinearity analysis of gene sequences between the *D. destructor* genome in this study and the published Dd-A1 genome.

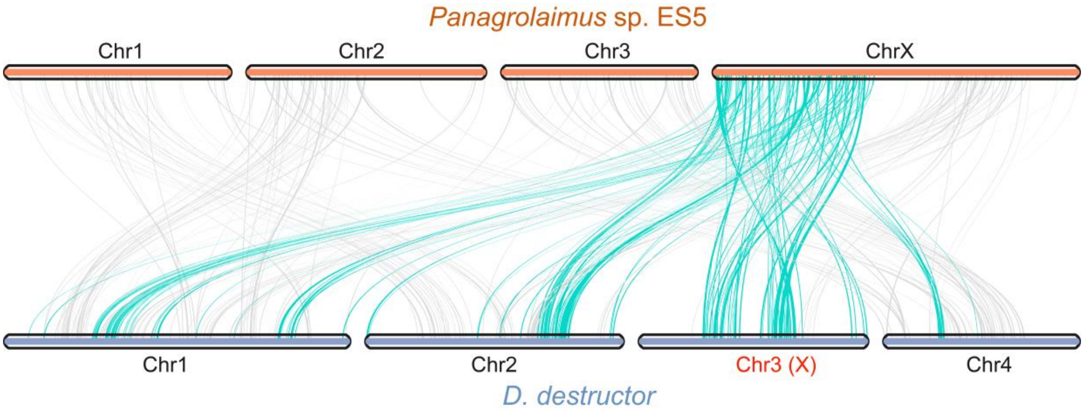


**Fig. S3.** Synteny analysis of protein-coding sequences between *D. destructor* and *Panagrolaimus* sp. ES5 suggests that chromosome 3 of *D. destructor* corresponds to its X chromosome.

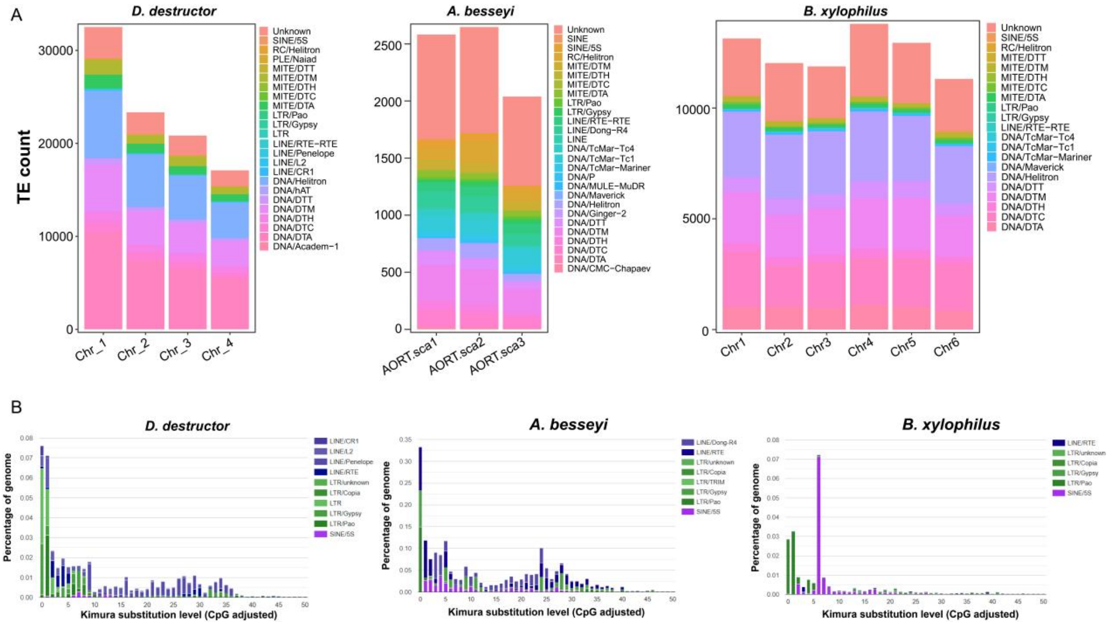


**Fig. S4.** The TE landscape of three migratory plant-parasitic nematodes. **A:** TE quantity and type statistics in *D. destructor*, *A. besseyi*, and *B. xylophilus*. B: The divergence of retrotransposon in three genomes. Kimura substitution levels were calculated for all retrotransposon TE copies found within each family to estimate the age of TE insertions. In the distribution, the closer the peak position is to 0, the younger the corresponding TE.

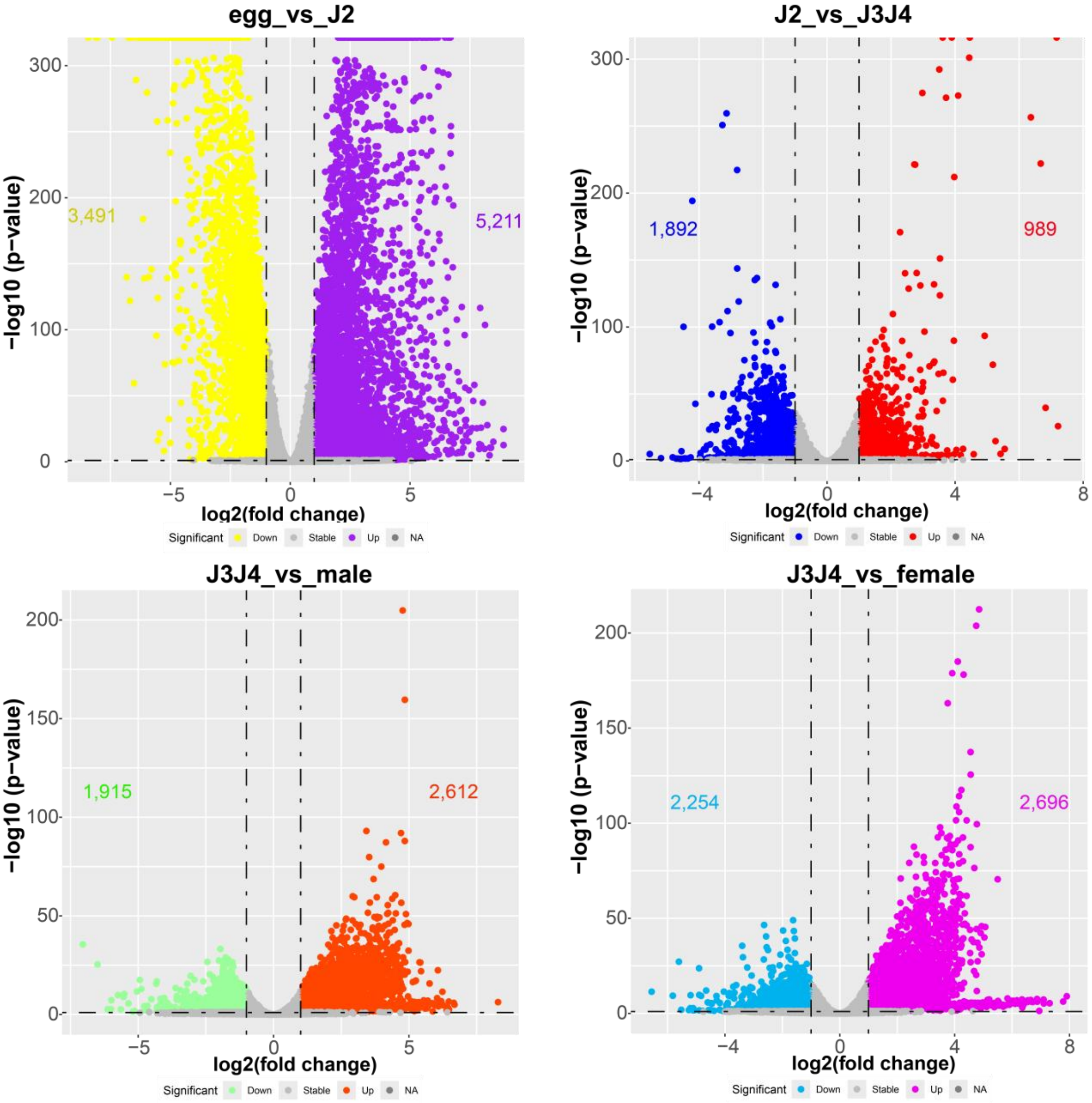


**Fig. S5.** Volcano plot showing differentially expressed genes between different developmental stages of *D. destructor*.

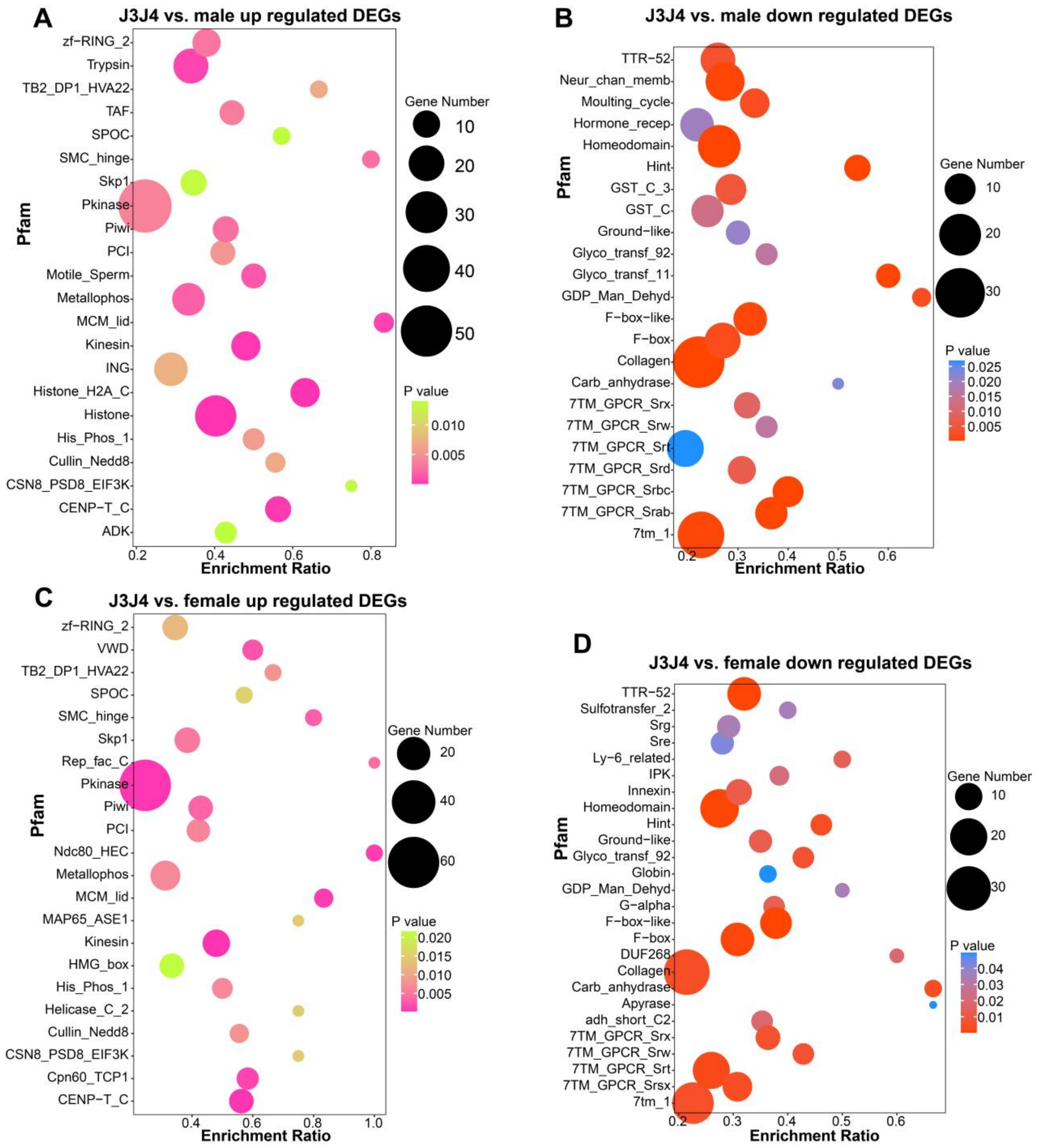


**Fig. S6.** Functional enrichment analysis of differentially expressed genes between J3J4 larvae and adults. **A:** Pfam conserved domain enrichment analysis of up-regulated DEGs between J3J4 and male. **B:** Pfam conserved domain enrichment analysis of down-regulated DEGs between J3J4 and male. **C:** Pfam conserved domain enrichment analysis of up-regulated DEGs between J3J4 and female. **D:** Pfam conserved domain enrichment analysis of down-regulated DEGs between J3J4 and female.

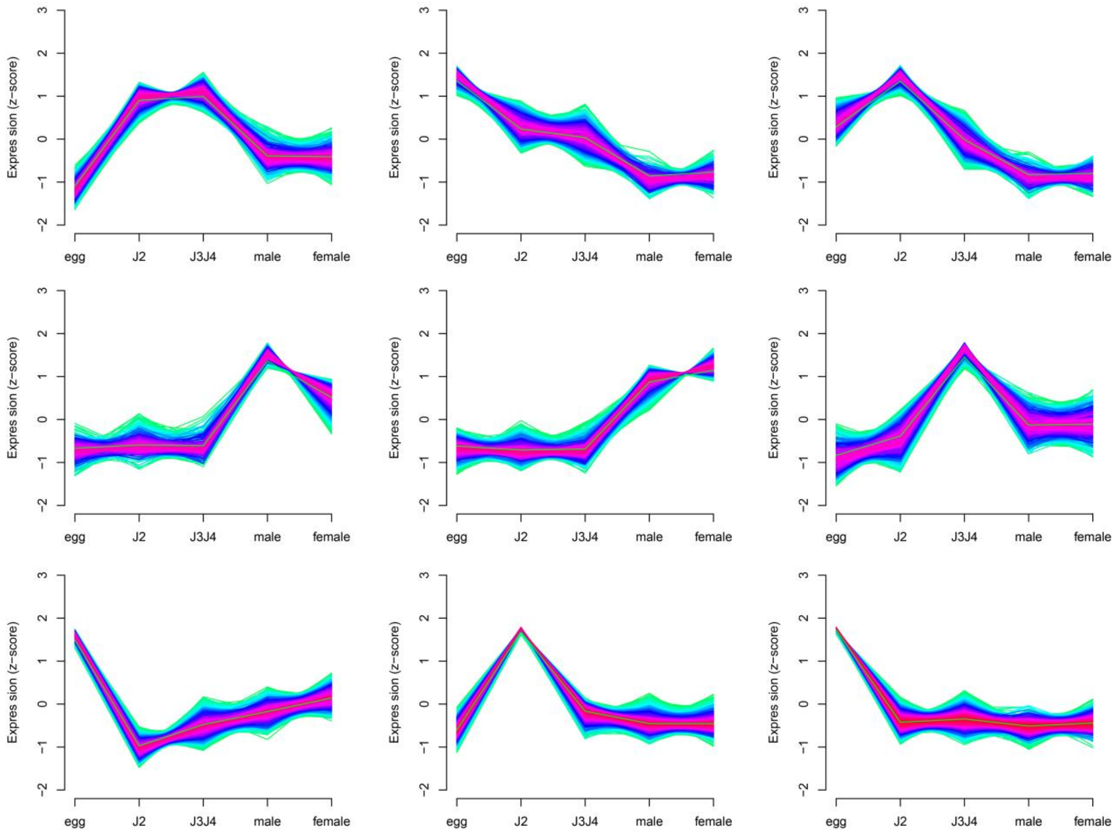


**Fig. S7.** Clustering of TPM values of gene expression between different stages of *D. destructor* shows age-specific expression profiles.

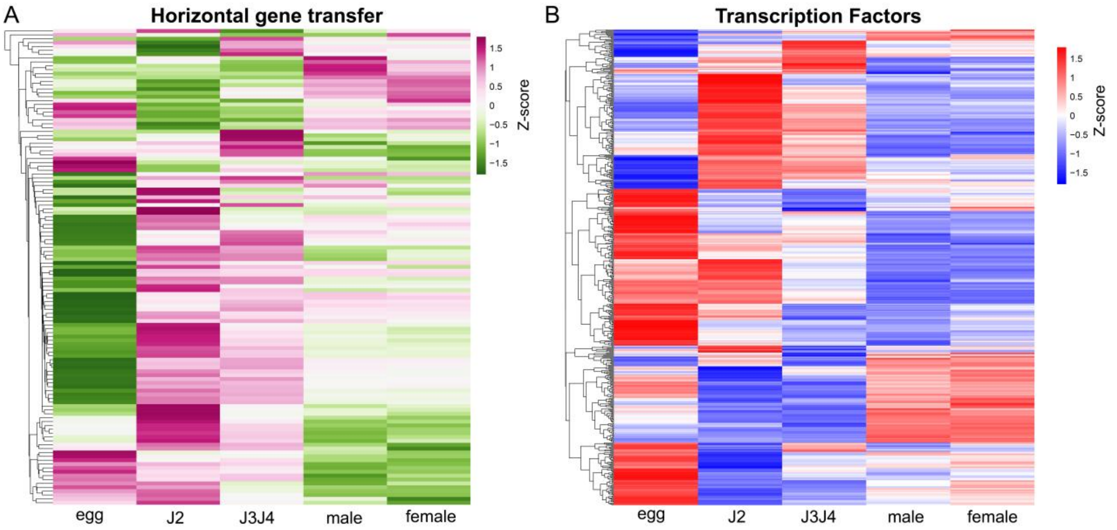


**Fig. S8.** The expression pheatmap of HGT and TF genes during the development stage. **A:** The expression pheatmap of HGT shows most of HGT have the highest expression level in J2 stage. **B:** The transcription factor expression pheatmap shows they mainly regulate the gene expression in egg and J2 stage.

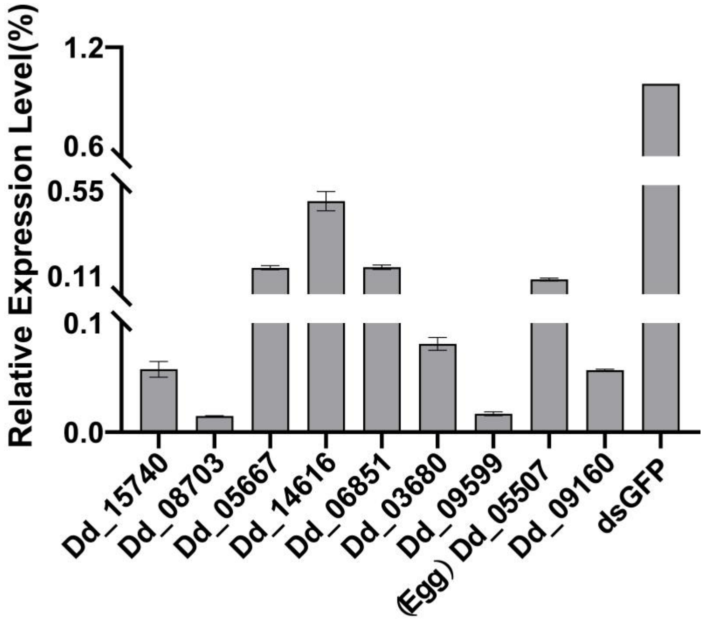


**Fig. S9.** qRT-PCR analysis of the relative expression levels of the candidate effector gene and transcription factor target genes following RNAi treatment.
