## Supplemental Tables for "Developmental dynamic transcriptomics reveals multiple effectors and transcription factors critical for *Ditylenchus destructor* parasitism"

**Table S1 The TE class summary of *D. destructor.***

| **Class** | **Count** | **bpMasked** | **%masked** |
| --- | --- | --- | --- |
| **DNA** | | | |
| Academ-1 | 195 | 36111 | 0.03% |
| DTA | 47915 | 8107832 | 7.73% |
| DTC | 5571 | 809809 | 0.77% |
| DTH | 6511 | 672334 | 0.64% |
| DTM | 26047 | 3408887 | 3.25% |
| DTT | 3288 | 358318 | 0.34% |
| Helitron | 32180 | 5769321 | 5.50% |
| hAT | 46 | 5104 | 0.00% |
| **LINE** | | | |
| CR1 | 21 | 8485 | 0.01% |
| L2 | 515 | 58173 | 0.06% |
| Penelope | 155 | 131728 | 0.13% |
| RTE-RTE | 40 | 47716 | 0.05% |
| **LTR** | | | |
| Copia | 55 | 74553 | 0.07% |
| Gypsy | 123 | 80132 | 0.08% |
| Pao | 41 | 55785 | 0.05% |
| unknown | 1 | 35 | 0.00% |
| **MITE** | | | |
| DTA | 6231 | 860565 | 0.82% |
| DTC | 12 | 2604 | 0.00% |
| DTH | 98 | 5753 | 0.01% |
| DTM | 7687 | 760357 | 0.72% |
| DTT | 797 | 64556 | 0.06% |
| **PLE** | | | |
| Naiad | 21 | 7002 | 0.01% |
| **RC** | | | |
| Helitron | 454 | 61234 | 0.06% |
| **other** | | | |
| SINE/5S | 78 | 13379 | 0.01% |
| Unknown | 15698 | 2619490 | 2.50% |
| **Total** | **153780** | **24019263** | **22.89%** |

**TableS2 The TE class summary of *B. xylophilus.***

| **Class** | **Count** | **bpMasked** | **%masked** |
| --- | --- | --- | --- |
| **DNA** | | | |
| DTA | 9843 | 1554461 | 1.99% |
| DTC | 31970 | 3822225 | 4.88% |
| DTH | 3556 | 586083 | 0.75% |
| DTM | 27556 | 3638249 | 4.65% |
| DTT | 6537 | 772902 | 0.99% |
| Helitron | 27382 | 3900335 | 4.98% |
| Maverick | 85 | 48089 | 0.06% |
| TcMar-Mariner | 398 | 76522 | 0.10% |
| TcMar-Tc1 | 312 | 87017 | 0.11% |
| TcMar-Tc4 | 149 | 42632 | 0.05% |
| **LINE** | | | |
| RTE-RTE | 20 | 5819 | 0.01% |
| **LTR** | | | |
| Copia | 20 | 1986 | 0.00% |
| Gypsy | 56 | 4413 | 0.01% |
| Pao | 44 | 67254 | 0.09% |
| unknown | 1 | 81 | 0.00% |
| **MITE** | | | |
| DTA | 1700 | 226910 | 0.29% |
| DTC | 1587 | 100198 | 0.13% |
| DTH | 50 | 2943 | 0.00% |
| DTM | 1953 | 276254 | 0.35% |
| DTT | 3 | 325 | 0.00% |
| **RC** | | | |
| Helitron | 162 | 36317 | 0.05% |
| **other** | | | |
| SINE/5S | 179 | 62343 | 0.08% |
| Unknown | 20951 | 5720422 | 7.30% |
| **Total** | **134514** | **21033780** | **26.87%** |

**Table S3 The TE class summary of *A. besseyi.***

| **Class** | **Count** | **bpMasked** | **%masked** |
| --- | --- | --- | --- |
| **DNA** | | | |
| CMC-Chapaev | 48 | 58437 | 0.13% |
| DTA | 16 | 13700 | 0.03% |
| DTC | 565 | 225108 | 0.48% |
| DTH | 349 | 101317 | 0.22% |
| DTM | 1356 | 528193 | 1.13% |
| DTT | 301 | 98034 | 0.21% |
| Ginger-2 | 34 | 17943 | 0.04% |
| Helitron | 428 | 159828 | 0.34% |
| MULE-MuDR | 28 | 7590 | 0.02% |
| Maverick | 18 | 16493 | 0.04% |
| P | 45 | 17756 | 0.04% |
| TcMar-Mariner | 242 | 127690 | 0.27% |
| TcMar-Tc1 | 593 | 269030 | 0.58% |
| TcMar-Tc4 | 159 | 50790 | 0.11% |
| **LINE** | | | |
| unknown | 43 | 13168 | 0.03% |
| Dong-R4 | 419 | 228725 | 0.49% |
| RTE-RTE | 287 | 270351 | 0.58% |
| **LTR** | | | |
| Copia | 19 | 1068 | 0.00% |
| Gypsy | 168 | 166124 | 0.36% |
| Pao | 97 | 91498 | 0.20% |
| TRIM | 1 | 41 | 0.00% |
| unknown | 1 | 61 | 0.00% |
| **MITE** | | | |
| DTA | 143 | 33551 | 0.07% |
| DTC | 35 | 6856 | 0.01% |
| DTH | 2 | 417 | 0.00% |
| DTM | 319 | 58858 | 0.13% |
| **RC** | | | |
| Helitron | 488 | 546006 | 1.17% |
| **other** | | | |
| SINE | 17 | 2024 | 0.00% |
| SINE/5S | 207 | 126635 | 0.27% |
| Unknown | 3515 | 1577676 | 3.38% |
| **Total** | **9943** | **4814968** | **10.31%** |

**Table S4 Primer sequence used in this study**

| Primer ID | Primer sequence |
| --- | --- |
| dsDd_15740.1-F | TAATACGACTCACTATAGGGAATTCCGAGTAGCGGT |
| dsDd_15740.1-R | TAATACGACTCACTATAGGGAGCTACTGGATTGTCTA |
| dsDd_08703.1-F | TAATACGACTCACTATAGGGAAGATGCTATTTTTCTCAA |
| dsDd_08703.1-R | TAATACGACTCACTATAGGGATGGTTTTTTGGTTGT |
| dsDd_09599.1-F | TAATACGACTCACTATAGGGAGGACCCATGATGTCC |
| dsDd_09599.1-R | TAATACGACTCACTATAGGGTATTGCCTTGATCTTCC |
| dsDd_05507.1-F | TAATACGACTCACTATAGGGCACAGAACAATGGCAA |
| dsDd_05507.1-R | TAATACGACTCACTATAGGGAATTCGAACTCCATT |
| dsDd_09160.1-F | TAATACGACTCACTATAGGGAGCAACTCAAAATTGCTGAA |
| dsDd_09160.1-R | TAATACGACTCACTATAGGGAAGAAGAGTGAGCTCGGAT |
| dsDd_05667.1-F | TAATACGACTCACTATAGGGTCCATTCCTATTTCCAG |
| dsDd_05667.1-R | TAATACGACTCACTATAGGGATTGTCTATAGGCTTGG |
| dsDd_14616.3-F | TAATACGACTCACTATAGGGCCACGCTCCAATAAAT |
| dsDd_14616.3-R | TAATACGACTCACTATAGGGATATACGCCCTGTGCT |
| dsDd_06851.1-F | TAATACGACTCACTATAGGGACCCGAAAATGACAG |
| dsDd_06851.1-R | TAATACGACTCACTATAGGGACCAAGTTGAATGTCC |
| dsDd_03680.1-F | TAATACGACTCACTATAGGGTACAAGTGTTTCGTGATTG |
| dsDd_03680.1-R | TAATACGACTCACTATAGGGTTTCACCTGTTTAGTGT |
| dsGFP-F | TAATACGACTCACTATAGGGAGTGCCATGCCCGAAGGTTA |
| dsGFP-R | TAATACGACTCACTATAGGGTCTGCTAGTTGAACGCTTCC |
| qRT15740.1-F | CGTCAAACGATGAAGTGATGAT |
| qRT15740.1-R | ATTTGCGTCCAAATCCTGGTCC |
| qRT08703.1-F | CTAATGCTTATTGTGCTTGCGG |
| qRT08703.1-R | CCCAATTTCGAGAATATGGATGG |
| qRT09599.1-F | AAGTAACGAGCAACAGGATGGT |
| qRT09599.1-R | GCCAGTTATTGACAGCTGACAT |
| qRT05507.1-F | CGATATCTTTGTGCATCGATCG |
| qRT05507.1-R | GATCTTCATTATTCGGTCTGCG |
| qRT09160.1-F | GGATTTATCACTGTGGACGACGA |
| qRT09160.1-R | CCTCCACGACGACTGTAATTAG |
| qRT05667.1-F | AACTCCACCGCCGTCTTTAAC |
| qRT05667.1-R | TCGCAAGATTCTATGGAATGTGC |
| qRT14616.3-F | ACCGATCCATCGGCTAACCTCA |
| qRT14616.3-R | GGAAAAGTACGGAGAGAAGTGG |
| qRT06851.1-F | TAGGTTCATTCCTAGGAGGCAC |
| qRT06851.1-R | AGATATGGTCGCAATCCTTGCA |
| qRT03680.1-F | AAGTGTTTCGTGATTGTCTTGC |
| qRT03680.1-R | AATAGTCCATACCTTGCACGCC |
| qRTtba-F | ACATTCTTCAGTGAGACGCAATC |
| qRTtba-R | ACCTTGGAGACCGTGACATT |
| qRTact-F | CGGAACGCAAGTACTCTGT |
| qRTact-R | GGTCCAGATTCGTCGTATTCC |
